# Loss of Tbx3 in mouse eye causes retinal angiogenesis defects reminiscent of human disease

**DOI:** 10.1101/2022.04.20.488944

**Authors:** M. Derbyshire, S. Akula, A. Wong, K. Rawlins, E. Voura, W.J. Brunken, M.E. Zuber, S. Fuhrmann, A.M. Moon, A.S Viczian

## Abstract

**Purpose:** In Retinopathy of Prematurity (ROP), infants often have incomplete vascularization, affecting the temporal region. A factor expressed in this region during retinal development is the T-box factor, *Tbx3*, which has not been studied in the mammalian eye. The purpose of this study was to determine if *Tbx3* is required during eye formation for retinal angiogenesis.

**Methods:** Conditional removal of *Tbx3* from both retinal progenitors and astrocytes was done using the optic cup-Cre recombinase driver, *BAC-Dkk3-CRE* and analyzed using standard immunohistochemical techniques.

**Results:** With *Tbx3* loss, the retinas were hypovascular, as seen in patients with ROP and Familial Exudative Vitreoretinopathy (FEVR). Retinal vasculature failed to form the stereotypic tri-layered plexus in the dorsal-temporal region. Astrocyte precursors were reduced in number and failed to form a lattice at the dorsal-temporal edge. We next examined retinal ganglion cells, as they have been shown to play a critical role in retinal angiogenesis. We found that melanopsin expression and Islet1/2-positive retinal ganglion cells were reduced in the dorsal half of the retina. In previous studies, loss of melanopsin has been linked to hyaloid artery persistence, which we also observed in the *Tbx3* cKO retina, as well as in infants with ROP or FEVR.

**Conclusions:** Together, these results show that TBX3 is required for normal mammalian eye formation for the first time. This potentially provides a new genetic model for retinal hypovascular diseases.

## Introduction

Retinal development requires the synchronized formation of neural and vascular tissue for visual function. The early mouse retina remains avascular until differentiation of retinal cells causes an increase in metabolic demand. The differentiated cells release proteins that stimulate the migration and proliferation of astrocytes from the optic nerve just prior to birth^1–3^ Astrocyte precursors cling onto the axons of retinal ganglion cells as they migrate into the retina. ^4,5^ Migrating astrocyte precursors then produce signals to attract endothelial cells into the retina, which in turn, produces proteins that cause the astrocytes to differentiate. Therefore, coordinated activity of neural, astrocytes and endothelial cells are essential for nascent blood vessels to begin forming a steady retinal blood supply. Disruption of signals from either the vascular cells or neurons halts angiogenesis throughout the retina, creating a hypovascular retina seen in Retinopathy of Prematurity (ROP) and Familial Exudative Retinopathy (FEVR). ^1^ Mutations in Wnt signaling pathway components, like the Wnt receptor Frizzled4, are linked to these syndromes ^6–9^, and are responsible for 50% of FEVR cases. ^1^ The genetic mutations causing the remaining half of the FEVR patients are unknown. ^1^

The signals, which drive creation of the retinal vasculature, function in concert with other signals that cause regression of the vascular source during embryonic development, the hyaloid artery. The hyaloid artery is a transient fetal vascular structure in many vertebrates. Its regression is triggered by signals from intrinsically photosensitive retinal ganglion cells (ipRGCs). ^10,11^ Failure of the hyaloid vessels to regress leads to persistent fetal vasculature (PFV), which leads to intraocular hemorrhage and impairment of vision. Despite decades of study, and the importance to human disease, a complete understanding the molecular regulation of retinal vascular development is still lacking.

In this study, we have identified a key factor, *Tbx3*, in cell types affecting retinal vascular development. In the frog, *Xenopus laevis, tbx3* is expressed in the eye field prior to optic vesicle formation where it plays an essential role early in eye formation. ^12–16^ Later in ocular development, *Tbx3* is found in the dorsal half of the optic cup in both frog and mouse. ^13,14,16–18^ Functional studies of the role of *Tbx3* in mammalian eye formation have not been done. An initial gene deletion of murine *Tbx3* did not study the eye ^19^ and produced only a hypomorphic deletion. ^20^ Thus, a conditional ablation strategy was developed ^20,21^ and used successfully to study the role of *Tbx3* in several other tissues, including vascular cells during kidney development. ^22–28^ We investigated the role of *Tbx3* in mammalian retinal development for the first time using a combination of expression studies and reverse genetics to probe function. To our surprise, we found that *Tbx3* is expressed not only in embryonic retinal progenitors, but also in retinal astrocyte precursors at P0. To investigate the effect of *Tbx3* loss-of-function on retinal development, we crossed mice bearing the *Tbx3* conditional allele ^20^ with a Cre driver, BAC-Dkk3-Cre, that functions in retinal progenitors, astrocytes, but not endothelial cells. ^29,30^ In the *Tbx3* conditional knockout mice, we observed that the hyaloid artery persisted, and the retina was hypovascular. We found fewer astrocytes in the *Tbx3* conditional knockout retina at P9, and by P30, the astrocytic lattice was malformed in the dorsal retina. With *Tbx3* loss, we also found thinner optic nerves and fewer intrinsically photosensitive retinal ganglion cells. Together, these results suggest that TBX3 is required for mammalian eye formation and that this conditional knockout could potentially be a new mouse model for hypovascular diseases.

## Methods

### Mice

Generation of transgenic mouse strains, BAC-Dickkopf3:cre (*BAC-Dkk3-Cre*) ^29^, *Tie2-Cre*,^31,32^ *Tbx3*^*mcm*^, ^24^ *Tbx3*-floxed and *Tbx3*^*GR*^,^20,21^ were maintained on a C57Bl/6J background. Tamoxifen treatment was done as previously described. ^33^ Cre activity was assessed using the Ai9 Cre reporter line (B6.Cg-*Gt(ROSA)26Sor*^*tm9(CAG-tdTomato)Hze*^/J; The Jackson Laboratory, stock#: 007909). We used sibling conditional knockout (*BAC-Dkk3-Cre*^*+*^;*Tbx3*^*ΔFl/ΔFl*^) and wild-type (*BAC-Dkk3-Cre*-negative;*Tbx3*^*Fl/+*^) mice. At least two litters were used for analysis with the number of mice (N) indicated. We observed no difference in levels of TBX3 protein in male and female samples by western blot (Fig. 1L) and thus, performed our analysis on a random mixture of sexes. This study was approved by the Upstate Institutional Animal Care and Use Committee and adheres to the ARVO Statement for the Use of Animals in Ophthalmic and Vision Research.

**Figure 1.**
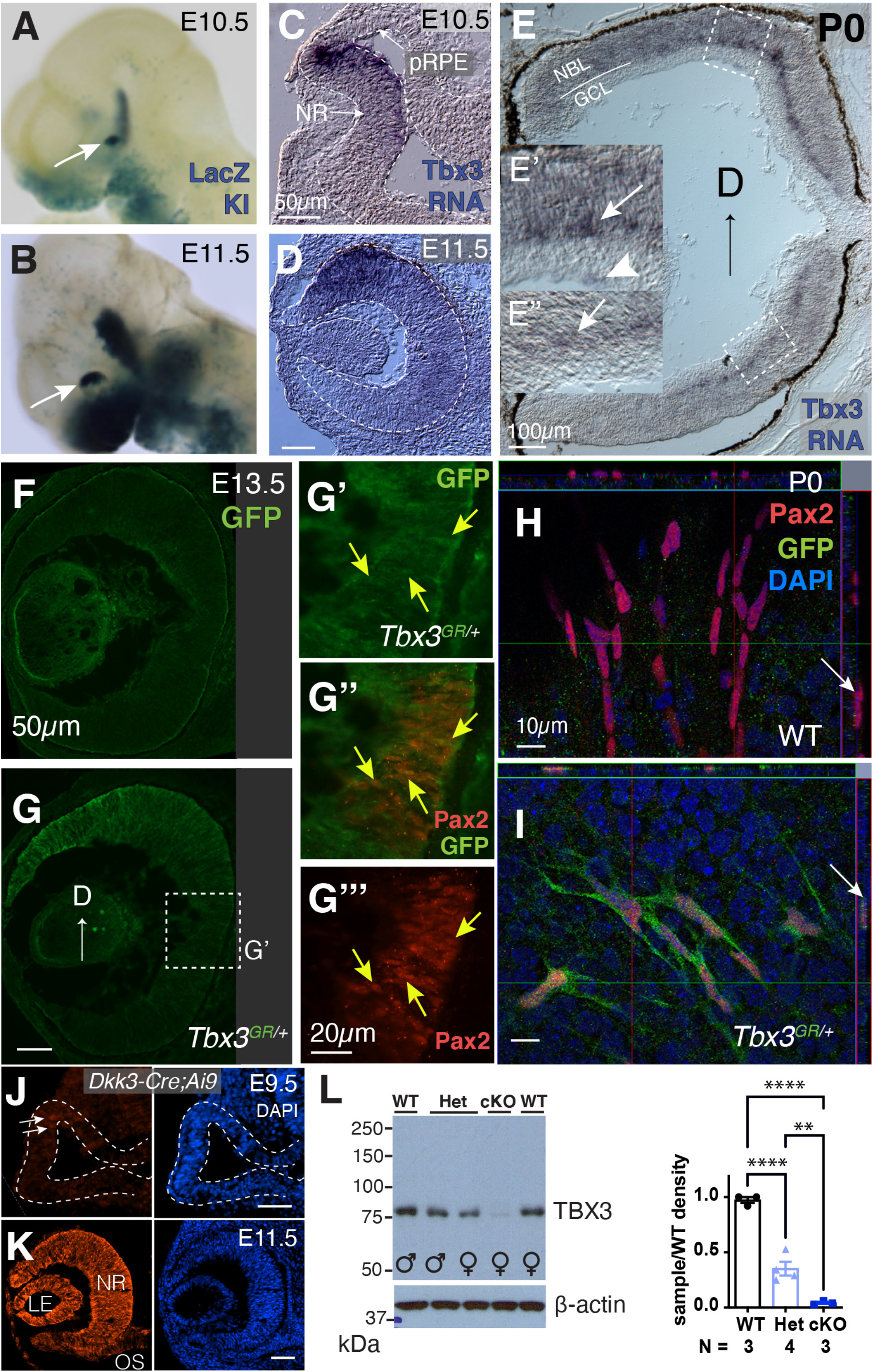
TBX3 expression in the neural retina and efficient conditional knock-out (cKO) using the *BAC-DKK3-Cre* driver. A,B) LacZ activity under the control of the endogenous*Tbx3*-locus (LacZ KI: *Tbx3*^*mcm*^). Arrow points to the dorsotemporal region of the eye at embryonic days 10.5 (E10.5) and 11.5 (E11.5). (C-D’) *In situ* hybridization using *Tbx3* antisense RNA at E10.5 and E11.5. Outline of developing eye in C,D; pRPE, presumptive RPE; NR, neural retina. (E) *Tbx3* mRNA expression in newborn retina. (E’,E”) Inlays of dorsal and ventral retinas. Arrows point to cells at the base of the neuroblast layer (NBL) and arrowhead to cells at inner limiting membrane adjacent to the ganglion cell layer (GCL). (F) Wild-type sibling of transgenic embryo (G) expressing GFP under the control of the endogenous *Tbx3*-locus (*Tbx3*^*GR/+*^) immunostained with anti-GFP antibody (green) at E13.5. Dashed box shows region magnified. (G’-G’’’) Close-up of boxed region in G immunostained with an anti-Pax2 (red) and GFP antibodies (see Fig. S1 for more images). (H) Confocal flat mount image of newborn retina from sibling wild-type and (I) *Tbx3*^*GR/+*^ mice immunostained with anti-Pax2 and GFP antibodies. Orthogonal view on top and side. Arrow points to cell that is positive for Pax2 in H and positive to Pax2 and GFP in I. (J,K) *BAC-Dkk3-CRE* expression pattern is detected using Ai9 reporter line expressing tdTomato. Cre activity is first detectable in the optic vessicle (dashed lines) at E9.5 (arrows) and is expressed throughout the neural retina (NR), lens epithelium (LE) and optic stalk (OS) by E11.5. (L) Western blot of P0 retinas collected from wild-type (WT, *Tbx3*^*Fl/+*^), heterozygous (Het, *BAC-Dkk3-CRE;Tbx3*^*ΔFl/+*^), and homozygous knockout (cKO, *BAC-Dkk3-CRE;Tbx3*^*ΔFl/ΔFl*^) siblings. Graph shows mean ± s.e.m. of 2 litters. No statistical differences in *Tbx3* expression were observed in males and females. Ordinary one-way ANOVA test with multiple comparisons was used to determine statistical significance. ****, p=0.00005; **, p≤0.01.

### Genotyping

Ear punches or tail snips (1-2 mm) were incubated overnight at 55°C in 100 µl lysis buffer (10mM Tris, pH 8.0, 100mM NaCl, 10mM EDTA, pH 8.0, 0.05% SDS) and 80 µg Proteinase K (Sigma; catalog #: P2308; 20 mg/ml stock). Samples were spun down at 14,000g for 15 minutes; 70µl supernatant was placed in 200µl ice cold isopropanol; DNA was pelleted at 4°C, 14,000g for 10 minutes and washed twice with 70% ethanol, dried and resuspended in 10 mM Tris, pH 8.0. PCR was performed as described in Table S1 with 0.25 units EconoTaq enzyme (Biosearch Technologies), 1X EconoTaq Buffer, 200µM dNTPs, 0.25µM primers for all reactions, except *Tbx3*^*GR*^ reactions that included 2% DMSO.

### Immunostaining

For sections, isolated eyes were fixed in 4% paraformaldehyde/1X PBS for either 30 min on ice (for *in situ* hybridization or GFP immunostaining) or overnight (for all other immunostaining). The tissue was washed in 1X PBS, and then, incubated with sucrose (5%, 10%, and 20%). The eyes were then mounted in O.C.T. (Tissue-Tek) and sectioned (20µm). We immunostained sections by using conditions that depended on the primary antibody (Table S2). Slides were washed 3 times in 1X PBST (1X PBS and 0.1% TritonX) and cover slipped with FluorSave (MilliporeSigma) and 2% DABCO (1,4-Diazabicyclo[2.2.2]octane; Sigma). Flat mounts were stained with isolectin B4 (IB4) and immunostained with antibodies, as described. ^34^ Reinforcement labels (Avery 5722) were attached to slides, retinas were placed inside each ring with mounting solution [FluorSave (MilliporeSigma) and 2% DABCO, and topped with a round cover slip.

### RNA in situ hybridization

RNA was extracted from mouse retinas using RNAzol RT (Molecular Research Center Inc.) and cDNA was generated using GOSCRIPT (Promega) reverse transcriptase according to the manufacture’s protocol. *Tbx3* cDNA was PCR amplified with Herculase II (Fisher) using the following primers: BamH1 mTbx3 Forward (5’ aat<u>ggatcca</u>CCATGAGCCTCTCCATGAG 3’) and Xho1 mTbx3 Reverse (5’ aatt<u>ctcgag</u>TTAAGGGGACCCGCTG 3’) according to the manufacture’s protocol. The PCR product was cut with BamH1 and Xho1 (New England Biolabs) and cloned directly into pCS2R cut with the same enzymes. DIG-labeled sense and antisense cRNA probes were manufactured from linear DNA made using these same two restriction enzymes. In situ hybridization was done as previously described. ^35,36^

### Microscopy

Images were taken using a Leica DM600B stereomicroscope outfitted with a MicroPublisher 6 camera (Teledyne Q-Imaging), and, where applicable, stitched together using the Volocity (v6.3) software. Confocal images were taken with the Zeiss LSM780 confocal and Zeiss Imaging Software. Images of WT and cKO samples were equally processed for brightness and contrast in Adobe Photoshop 2022. Cell counts and analysis were performed under double-blind conditions. Sample types were revealed when entering numbers into the statistics software (Prism v9.3).

### Measurement of vasculature

We measured the surface area occupied by retinal structures and the volume using Volocity software. The ratio of retinal structure to whole retinal surface was calculated and statistically analyzed. The migration of Pax2+ cells was measured within 500µm of the dorsal petal (counting 10 cells per sample). The area occupied by Pax2+ cells was measured by adjusting the threshold in ImageJ and comparing that to the area of the dorsal retina.

### Western immunoblotting

Protein was extracted from both retinas of sibling newborns and processed for western blot analysis, as previously described ^37^ Anti-TBX3 antibody (1:2000; Bethyl Lab, Inc.) in 5% milk, 0.1% Tween/1XTBS was used.

### Statical analysis

Prism (v9.3) was used for statistical analysis. Measurements were done blind to the genotype of the sample. We normalized data by taking the average wild-type measurement in each litter and dividing it into each sample in that litter. To determine the density of OPN4-positive cells, we divided retinal petal surface area (Volocity) into the total cells counted. The average density for WT samples was calculated; this number was used to divide into the density for all sibling samples. Graphs and data are reported as mean ± the standard error of the mean (s.e.m.). Each point on a graph is the average of both eyes from a single animal (N).

## Results

To determine the expression pattern of *Tbx3* in the embryonic eye, a tamoxifen-inducible Cre was knocked into the *Tbx3* gene locus (*Tbx3*^*mcm/+*^) and then crossed with the *Rosa*^*LacZ*^ reporter line (*Tbx3*^*mcm/+*^; *Rosa*^*LacZ/+*^; Fig. 1A, B). ^24^ Consistent with earlier studies, *LacZ* was expressed at E10.5 and E11.5 in the dorsotemporal region of the optic cup with gradual reduction in the ventral eye. ^17,38^ *In situ* hybridization to detect *Tbx3* mRNA showed a similar gradient of transcript expression (Fig. 1C,D). In newborn mice, we find that *Tbx3* mRNA is confined to the base of the neuroblast layer (arrows, Fig. 1E-E’’) and in cells at the inner limiting membrane (arrowhead, Fig. 1E’). To better visualize TBX3 expression, we used the recombined *Tbx3*^*GFP*^ conditional reporter allele, *Tbx3*^*GR/+*^, in which GFP has been knocked into the endogenous *Tbx3* gene (TBX3-GFP). ^21^ Relatively little GFP staining was detected in wild-type sibling embryos (Fig. 1F), while in*Tbx3*^*GR/+*^ embryos, we detected embryonic TBX3-GFP in the same gradient pattern with a few GFP-positive cells at the optic nerve head (arrows, Fig. 1G’). At E13.5, Pax2 is expressed in the optic nerve head in cells fated toward an astroglial lineage. ^3,39,40^ We detected GFP at low levels co-staining in some Pax2-positive cells (arrows, Fig. 1G’-G’’’ and Fig. S1A-B’’) with Pearson’s correlation co-efficient for GFP/Pax2-positive retinal cells averaged 0.337 ± 0.05 (>4 sections measured per eye; N=2), indicating low correlation. At birth, Pax2 marks astrocyte precursors as they migrate into the retina. ^41,42^ We flat mounted retinas at P0, immunostained with anti-Pax2 and GFP antibodies, and imaged whole retinas (Fig. S1 C,D and Fig. 1H,I). Flat mounts of P0 retinas from sibling WT have no visible GFP expression, yet TBX3-GFP mice show GFP/Pax2 co-labeling (Fig. 1H,I and Movies 1,2). Together, these data suggest that *Tbx3* is initially expressed in retinal progenitors, and possibly some Pax2-positive glial progenitors while later, it is expressed in astrocyte precursors, as well as cells in the neuroblast layer.

To investigate the effects of *Tbx3* loss on early eye formation, we selected the *BAC-Dkk3-CRE* mouse line. ^29^ To confirm its specificity, *BAC-Dkk3-CRE* mice were crossed with the Ai9 reporter line that expresses tdTomato in the presence of CRE activity. As previously reported, we found weak expression of the tdTomato reporter in E9.5 optic vesicles (arrows, Fig. 1J) that was detected throughout the neural retina, optic stalk and in the lens epithelium by E11.5 (Fig. 1K and S1E-E’’), but not in endothelial cells (dashed arrow, Fig. S1E’’). In *BAC-Dkk3-CRE* postnatal retinas, there was a gap in tdTomato expression in blood vessels, but not the surrounding neurons and astrocytes (arrow, Fig. S1F-F’’’), confirming that Cre activity was absent in these retinal endothelial cells. To ensure that our protocol detected tdTomato in blood vessels, we used *Tie2-Cre;Ai9* retinal flat mounts stained with the same blood vessel markers, which perfectly matched the expected vascular structures (Fig. S1G-G’’’). Together, these results suggest that *BAC-Dkk3-CRE* drives Cre recombinase in neurons, astrocytes, but not endothelial cells. Western blot analysis confirmed the loss of TBX3 protein in the *BAC-Dkk3-CRE;Tbx3*^*ΔFl/ΔFl*^ retina (Fig. 1L). Preliminary observations indicated abnormal blood vessel formation in the eyes of *BAC-Dkk3-CRE;Tbx3*^*ΔFl/ΔFl*^ mice. Therefore, we investigated the effect of *Tbx3* loss on retinal vasculature development.

Retinal vasculature develops postnatally in mice and blood vessels reach the edge of the retina by P9. ^43,44^ Therefore, to determine the effect of *Tbx3* loss on angiogenesis, we stained P9 retinal flat mounts using isolectin B4 (IB4) to visualize vascular endothelial cells (Fig. 2A,A’). *BAC-Dkk3-CRE;Tbx3*^*ΔFl/ΔFl*^ mice (*Tbx3* cKO) had severe disruption of vasculature development, in that the blood vessels failed to advance to the edges of the retina (Fig. 2B,B’). The hyaloid artery (HA) regresses postnatally and is substantially reduced by P8. ^11^ While we observed remnants of this structure in some wild-type samples (arrow, Fig. 2A’), the vessel were easily observed in *Tbx3* cKO retinas (Fig. 2B,B’). Measuring retinal surface areas covered by the vasculature (Fig. 2C), we found a decrease in the ventral vascular coverage in comparing *Tbx3* cKO and wild-type siblings (71 ± 5.9%), yet the dorsal *Tbx3* cKO retina exhibited more loss (Fig.2C; 24.9 ± 3.0%). These results indicate a requirement for *Tbx3* in the normal development or maintenance of the early postnatal retina vasculature. It is also possible that *Tbx3* loss simply causes a delay in forming the retinal vasculature, so we next looked at blood vessels after they matured.

**Figure 2.**
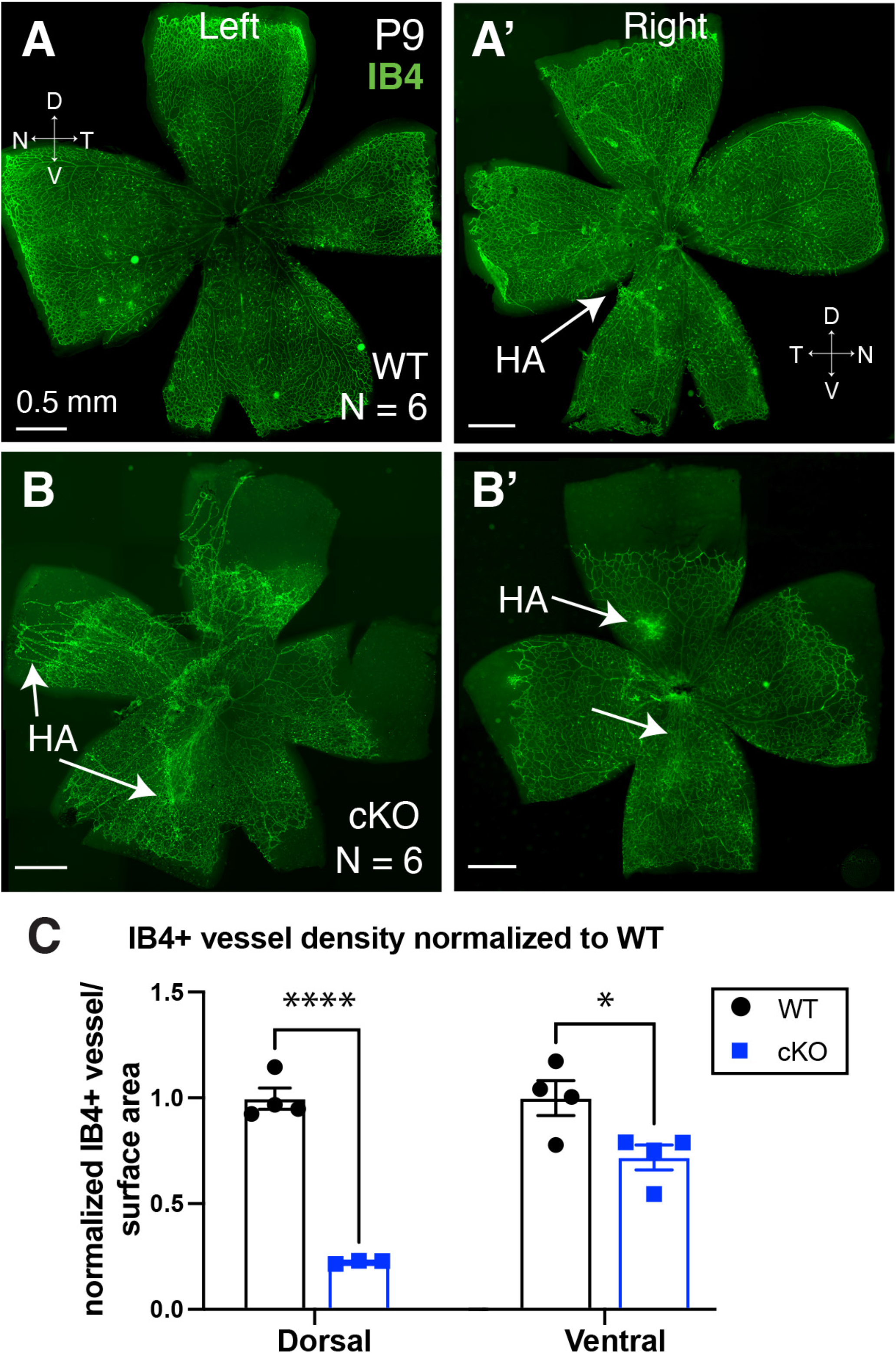
Tbx3 loss results in abnormal blood vessel development. (A,A’) Flat mount P9 retinas from wild-type (WT) and (B,B’) conditional *Tbx3* knockout retinas (cKO) stained with isolectin GS-B4 (IB4, green) in left and right eyes. Arrows indicate location of hyaloid artery (HA). C) Ratio of the vascularized to total retinal surface area in the dorsal and ventral retina normalized to average WT in each litter. A 2way ANOVA with Šidák’s multiple comparisons test was used; ****, p ≤0.0001; *, p=0.01.

Retinas were collected from P26 mice to determine the effect of *Tbx3* loss on the mature vascular plexus. In contrast to the wild-type structure (Fig. 3A-C), the vascular plexus of *Tbx3* cKO retinas was abnormal (Fig. 3D-F). Confocal optical sections showed the superficial layer did not extend to the dorsotemporal peripheral retina in *Tbx3* cKO mice (Fig. 3D,F, dashed red line marks areas; Movie 3). In addition, the dorsal vein was thinner and missing from the edge of conditional knockout retinas (red arrowheads, Fig. 3G vs. 3H; Movie 3). The *Tbx3* cKO vascular plexus also appeared less complex when compared to wild-type siblings (Fig. 3G,H). To further define the apparent reduction in complexity, we measured both the vasculature’s branch point and length density in *Tbx3* cKO and wild-type animals (Fig. 3I; quantitation shown in Fig. S2). We found a reduction in both branch point density (WT, 1537 ± 99; cKO, 1034 ± 106 junctions/mm^2^) and vascular length density (WT, 6.48 ± 0.12; cKO, 5.20 ± 0.26 mm/mm^2^) in mutants (Fig. 3I). Confocal imaging of central retinal sections from controls showed the normal tri-layered vasculature and inter-plexus branching (Fig. 3J and Movie 4). In contrast, *Tbx3* cKO retinas lacked the superficial plexus (Fig. 3K and Movie 5). Cryostat sections show vessels in all three layers in the wild-type (Fig. 3L and Fig. S3A), while in the absence of *Tbx3*, some regions (as in dashed regions shown in Fig. 3F) were missing both the superficial and deep layers (Fig. 3M and Fig. S3B). Glial fibrillary acidic protein (GFAP) labels Müller glia and astrocytes at this age. We detected less dense GFAP staining in the *Tbx3* cKO retina and that the outer nuclear layer is thinner at the dorsal edge in the mutants (bracket, Fig. 3M). Less GFAP staining suggests fewer astrocytes in the *Tbx3* cKO retina. Together, these results indicate that the formation of a normal adult retinal vasculature requires *Tbx3*.

**Figure 3.**
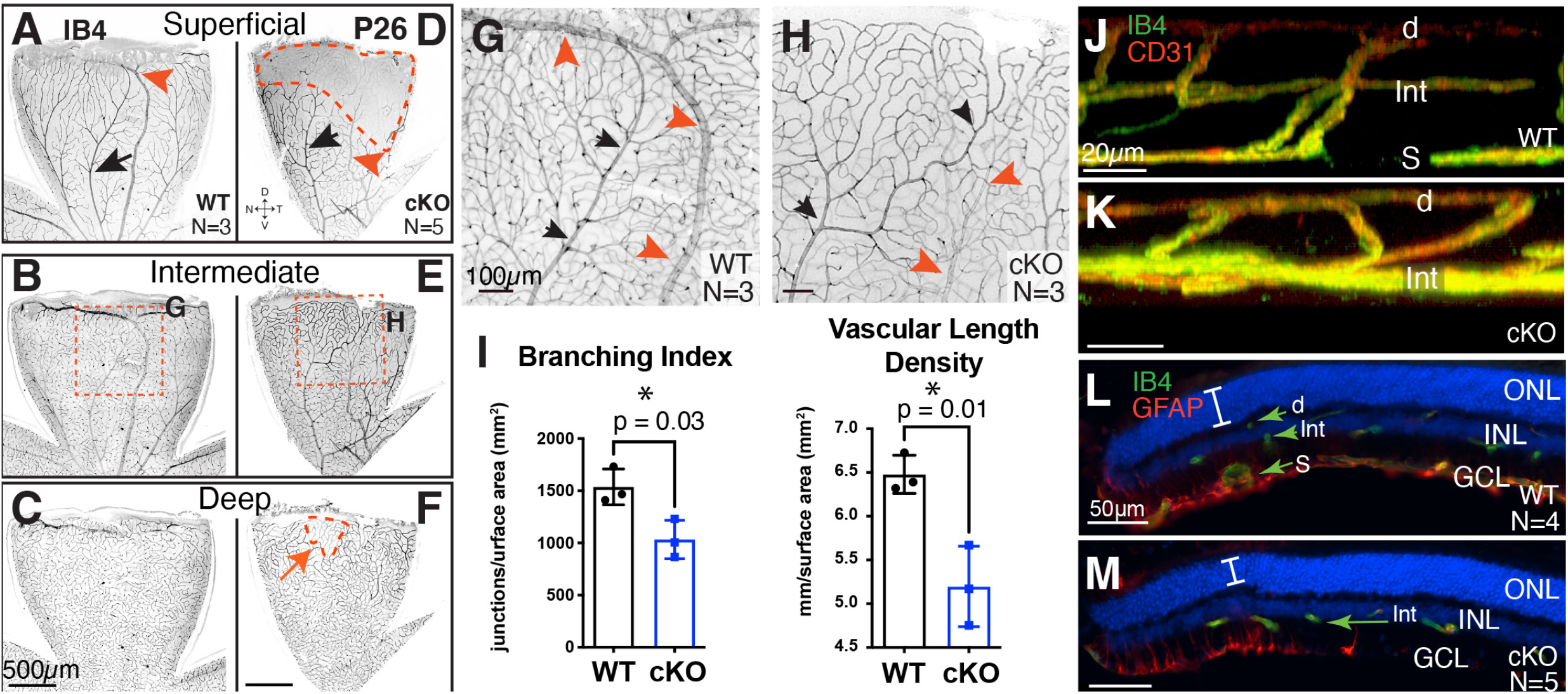
*Tbx3* cKO retina fails to form superficial plexus in the adult dorsal retina. (A-F) Confocal flat mount images of P26 dorsal retinas from wild-type (WT; A,B,C) and conditional knockout (cKO; D,E,F) mice stained with isolectin GS-B4 (IB4). Confocal images are inverted. The superficial (A,D), intermediate (B,E) and deep (C,F) plexus from the same retinal flat mounts are shown. (G,H) Maximal projection images from a stereomicroscope simultaneously showing all three plexi from the region indicated in B and E. Artery (black arrows) and vein (red arrowheads) in each panel are marked. (I) Branching index (branch points/retinal surface area) and vascular length density (vessel length/retinal surface area) from maximum projections images (see Fig. S2 for quantitation). Statistical significance was determined using an unpaired, two-tailed T-test; p-values on graph. (J,K) Still 3D rendered confocal images of retinal plexus in dorsal retina of WT (J) and cKO (K) stained with IB4 (green) and anti-CD31 antibody (red). (L,M) Transverse cryostat sections of P26 retina stained with IB4 (green) and anti-GFAP antibody (red) to mark endothelial cells and astrocytes, respectively. Superficial (S), intermediate (Int) and deep (d) vascular plexus are marked; outer nuclear layer (ONL), inner nuclear layer (INL), and ganglion cell layer (GCL) can be seen with nuclei marker, DAPI (blue). Note that the anti-GFAP (raised in mouse) antibody lightly stains vascular cells, as these mice were not perfused.

If the reduction in angiogenesis in the *Tbx3* cKO begins at this early time point, then we would expect the lattice to be disrupted in mutant mice. Flat mounts at P2 showed reduced endothelial cells and astrocytes in the *Tbx3* cKO retina (Fig. 4B,B’). Retinal astrocyte precursors express the paired-box transcription factor, Pax2, and the growth factor receptor, PDGFRα, and, as they mature express not glial fibrillary acidic protein (GFAP). ^41,42^ We found stunted migration and fewer Pax2-positive astrocyte precursors in the dorsal and ventral retinas at P2 (Fig. 4C-F). As we found TBX3 expressed in early astrocyte precursors (Fig. 1I), the stunted growth or migration of astrocyte precursors suggests that TBX3 may be required by these cells as they move into the retina. We further reasoned that lattice formation may recover if TBX3 is only required at this initial stage, and therefore, next looked at older animals.

**Figure 4.**
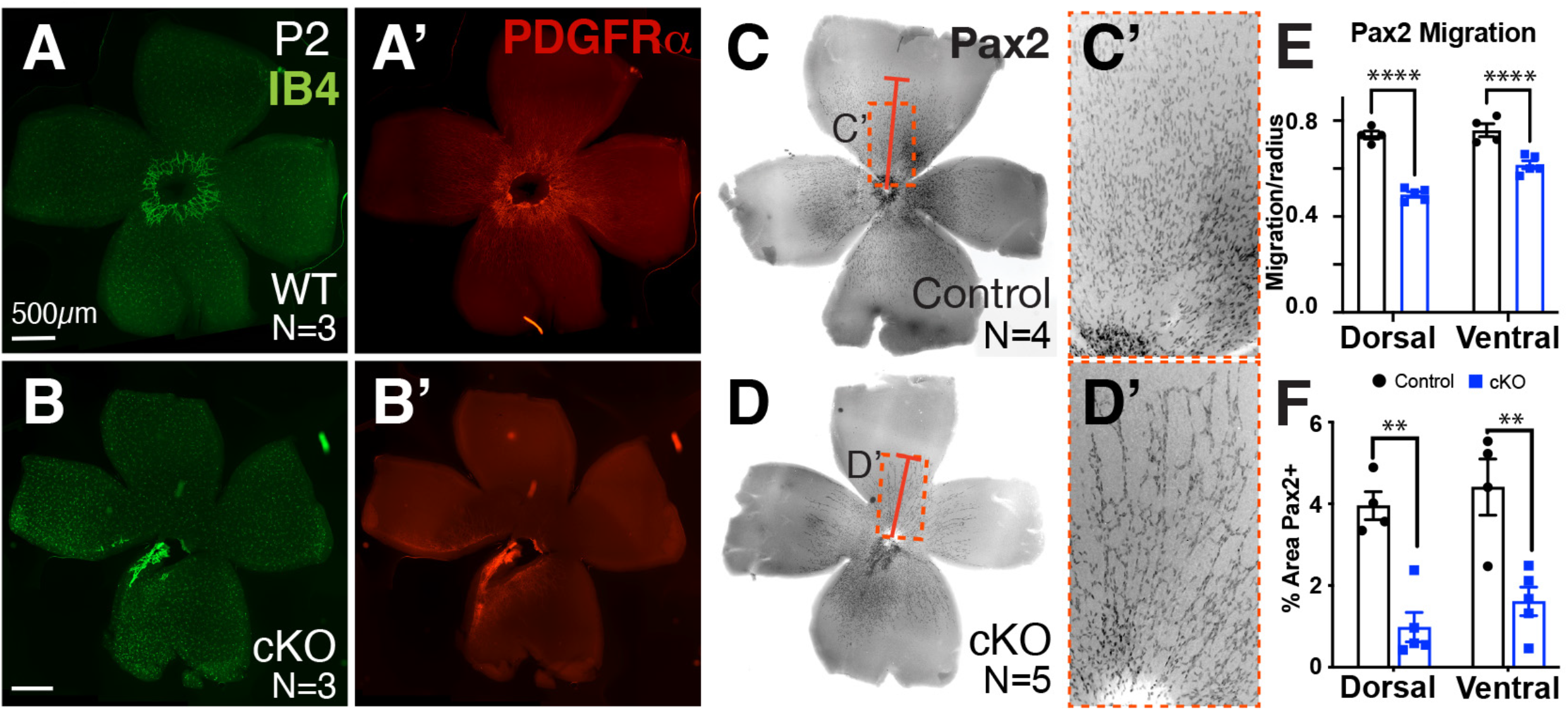
Fewer astrocyte precursors in *Tbx3* cKO retinas, starting as early as P2. (A,B) Endothelial cells are marked with IB4 (green) in WT and *Tbx3* cKO flat mounted retinas at P2. (A’,B’) Same samples immunostained for astrocyte precursors using anti-PDGF receptor alpha (PDGFRα) antibody. (C,D) Inverse images of flat mount retinas from P2 WT and *Tbx3* cKO stained with a second astrocyte precursor marker, anti-Pax2 antibody (black). Red bar marks Pax2-positive cells furthest away from the optic nerve head in each panel. Dashed yellow box indicates images magnified in panels C’,D’. (E) Quantitation of Pax2-positive cells furthest away from optic nerve head in heterozygote (control, N = 4) and conditional knockout (cKO, N = 5) retinas. Distance was averaged measure of 10 cells per dorsal and another 10, ventral. Statistical significance was determined using a 2way ANOVA with Tukey multiple comparisons test; ****, p <0.0001. (F) Graph showing quantitation of the percentage of surface area occupied by dorsal and ventral retinas in control (grey) and cKO (white); statistical significance was determined using a 2way ANOVA with Tukey multiple comparisons test; **, p <0.01; N, number of mice.

By P9, astrocytes reached the edge of the wild-type retina (Fig. 5A), yet failed to do so in the conditional knockout (Fig. 5B). Even at one month of age, astrocytes failed to reach the dorsal edge of the cKO retinas (compare Fig. 5C to 5D,D’). Double labeling astrocytes and endothelial cells, we found an abnormal vascular plexus in areas without a stereotypical astrocyte lattice (Fig. 5D’-D’’’). We compared the surface area occupied by astrocytes at P9 and P30, normalized to the average wild-type retinal surface area. We observed a statistically significant difference between wild-type and *Tbx3* cKO retinas at P9 and P30 (P9 WT, 1.0 ± 0.01; P9 cKO, 0.58 ±0.02; P30 WT, 0.97±0.01; P30 cKO, 0.76 ±0.03; Fig. 5E). We also observed significant growth between P9 and P30 (p<0.001), indicating that signals promoting astrocyte lattice formation persist in the absence of TBX3 (Fig. 5E). These results indicate that astrocytes require TBX3 to form the astrocytic lattice required for normal retinal vasculature formation. Astrocyte lattice formation can also be disrupted with the loss of retinal ganglion cells, as in the *Atoh7* knockout retina. These mice grow a severely stunted astrocytic lattice. ^4,45^ Thus, it is possible that TBX3 is indirectly affecting astrocytes by functioning within ganglion cells. Previous studies have also shown that hyaloid artery regression is dependent on signals from intrinsically photosensitive retinal ganglion cells. ^10,11^ We observed that the hyaloid artery persisted in the mutant retina (Figs. 2B,B’ and 5B). Therefore, we next investigated whether retinal ganglion cell formation requires TBX3.

**Figure 5.**
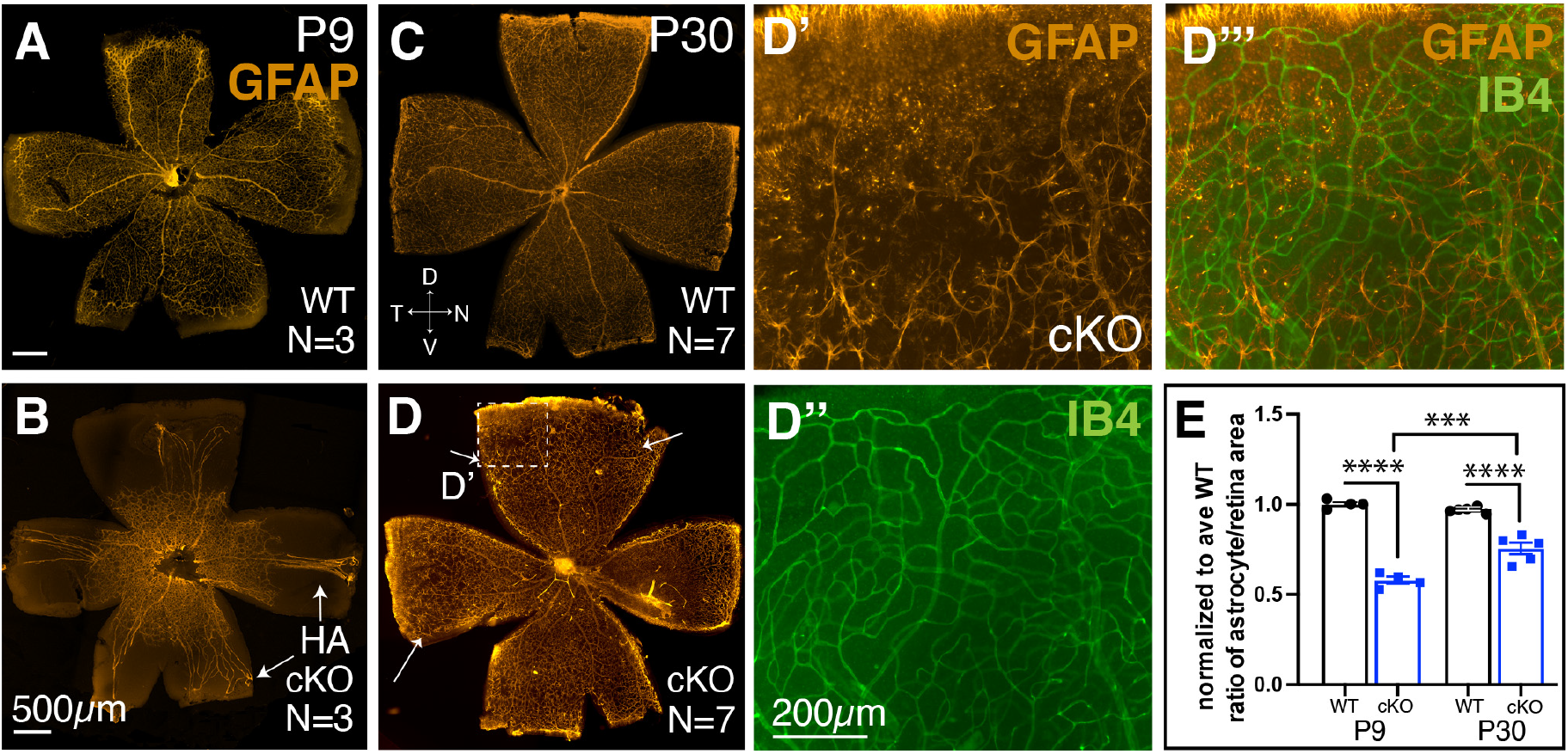
Astrocyte lattice fails to form in dorsal-temporal *Tbx3* cKO retina. (A,B) Flat mount P9 retinas immunostained with mature astrocyte marker, anti-glial fibrillary acidic protein (GFAP) antibody. Arrows point to the hyaloid artery (HA). (C,D) Flat mount P30 retinas from WT and cKO mice. (D) Arrows indicate lattice thinning. (D’-D’’’) Magnified view of boxed region in (D) shows dorsal region where astrocyte lattice fails to reach the retinal edge. (D’) Astrocytes, GFAP (orange); (D’’) endothelial cells, IB4 (green); and (D’’’) overlay. (E) Graph quantifying the normalized ratio of astrocyte occupation compared to the retinal surface area for WT and cKO retinas. Dorsal is up in all panels; statistical significance was determined using an ordinary 2way ANOVA with Tukey multiple comparisons; ****, p≤0.000.1; ***, p ≤0.001; N, number of mice.

We found that optic nerves were significantly thinner in cKO mice compared to their WT littermates (Fig. 6A-C), measuring 71 ± 3.9% of the average WT optic nerve (Fig. 6D). Using the anti-Islet1/2 antibody, we observed a significant decrease in dorsal Islet1/2-positive retinal ganglion cells (RGCs) in the cKO retina, but no difference in the ventral region (dorsal WT: 37.7 ± 2.8 cells; dorsal cKO: 27.7 ± 2.4 cells; Fig. 6E-G). This suggests that *Tbx3* is required by a subset of Islet1/2-expressing dorsal RGCs. Islet1 is thought to regulate the formation of most RGCs, including melanopsin-expressing (OPN4) intrinsically-photosensitive ganglion cells (ipRGCs). ^46^ We stained flat mounts with the anti-OPN4 antibody and measured the density of OPN4-positive cells for each sample, normalized to the average wild-type control. When blood vessels were stunted at P9, we observed significant reduction of ipRGC density in the dorsal (WT: 1.00 ± 0.11; cKO: 0.57 ± 0.06) and nasal/temporal petals (WT: 0.83 ± 0.07; cKO: 0.46 ± 0.06), but not in the ventral petals (Fig. 6H-J). Overall, we found significantly less ipRGCs in cKO retinas at P9 (WT: 1.00 ± 0.10; cKO: 0.63 ± 0.09). This difference persists when measured at P30 (WT: 1.00 ± 0.09; cKO: 0.68 ± 0.02), suggesting that the reduction in ipRGCs was not due to a delay of differentiation (Fig. 6K-L). Together, these results indicate that TBX3 is required for a subset of OPN4-expressing ipRGCs in the dorsal half of the retina, which control vascular formation in this region.

**Figure 6.**
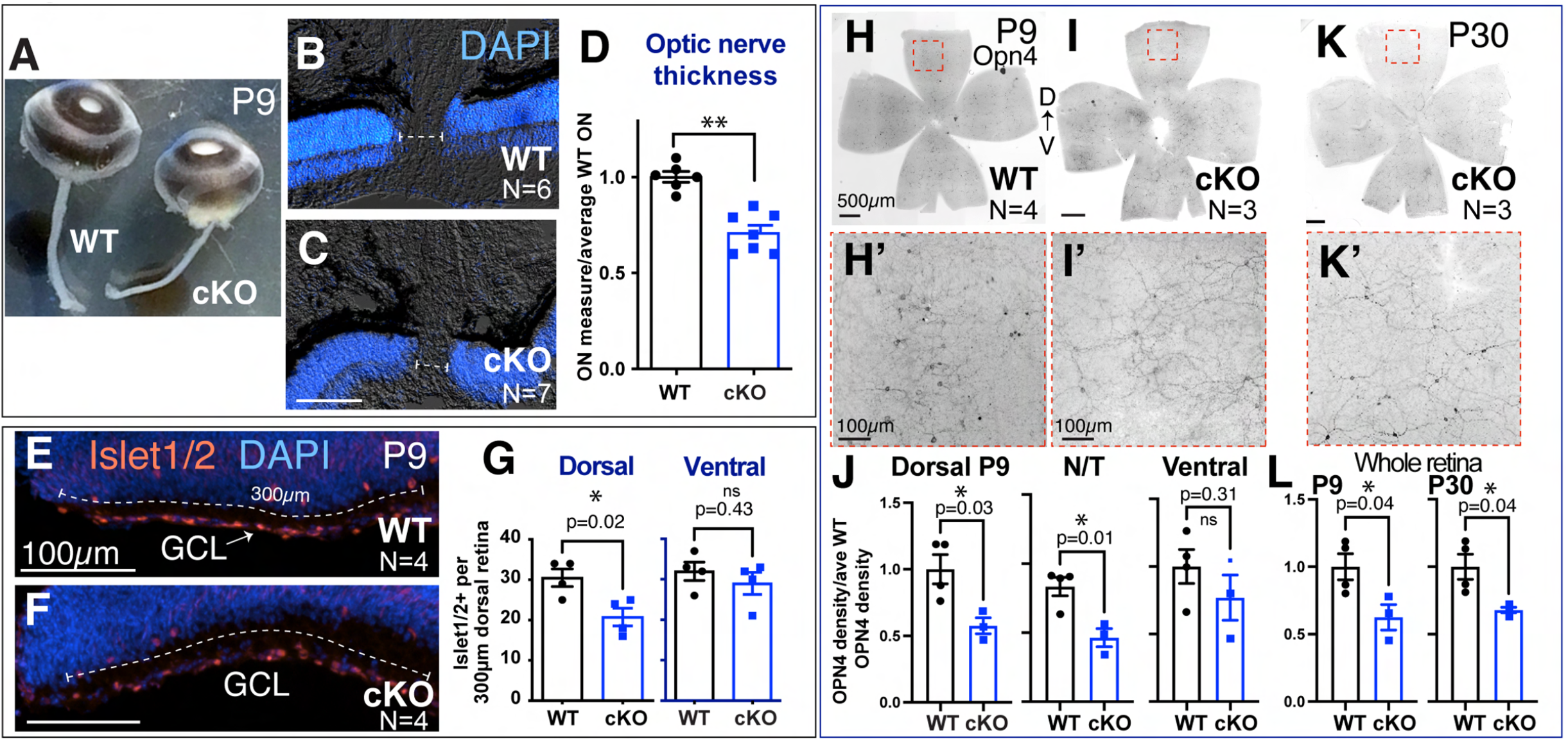
Dorsal specific loss of RGCs with loss of *Tbx3*. (A) Snapshot of whole eyes with optic nerves from P9 WT and cKO siblings. (B,C) Sections of WT and cKO eyes stained with nuclei marker (DAPI) to visualize neural retina. Dashed line indicates optic nerve thickness in each sample. (D) Graph quantitating the optic nerve thickness in WT and cKO eyes. Statistical significance was measured using an unpaired, two-tailed T-test; **, p = 0.005. (E-F) Transverse cryostat sections of P9 WT and cKO dorsal tip immunostained with anti-Islet1/2 antibody. The number of Islet1/2-positive cells in the first 300μm of the most dorsal and ventral retina were counted (dashed line). (G) Graphs of the Islet1/2-positive cells in the dorsal and ventral regions. (H-I’) Inverse images of P9 flat mount WT and cKO retinas immunostained with anti-OPN4 antibody marking intrinsically photosensitive ganglion cells (ipRGCs). Dashed box shows area magnified in H’ and I’. Density of cells were measured in dorsal, nasal/temporal (N/T) and ventral petals and normalized to WT siblings. (J) Graphs showing quantitation of OPN4-positive cells in each region. (K) Inverse image of P30 cKO flat mount with magnified area (dashed box) shown in panel K’. (L) Graphs of OPN4-positive cells per whole retina of P9 and P30 samples (WT, N = 4; cKO, N=3). An unpaired, two tailed T-test were used to determine statistical significance; p-values on graphs in panels G,J,L; N, number of mice.

## Discussion

Our results provide the first evidence that TBX3 is required for mammalian eye formation. We have found that *Tbx3* mRNA and protein is expressed in embryonic retinal progenitors and postnatal astrocyte precursor cells. When we remove *Tbx3* from the embryonic optic cup, the hyaloid artery persists in P9 mice and retinal vascular formation is stunted. At P9, the vascular defect is found throughout the retina, but more severe in the dorsal region. The dorsal vascular plexus continues to be affected by *Tbx3* loss in the older animals, including the loss of the superficial layer of the vascular plexus and the astrocyte lattice. We observe that in the absence of *Tbx3*, fewer astrocyte precursors are present in the retina. As the retina matures, astrocyte lattice formation recovers in all but the dorsal region. To our knowledge, localization of the defect to the dorsal intraretinal vasculature is the first report of a spatially distinct defect in this vascular bed. In contrast, loss of Aldh1a1, also expressed in dorsal retinal neurons affects the dorsal choroidal, or outer retinal vasculature. ^47^ In the dorsal neural retina area with *Tbx3* conditionally removed, we find fewer Islet1/2-positive and melanopsin-expressing intrinsically photosensitive retinal ganglion cells. One interpretation is that these neurons fail to attract or stimulate the proliferation and/or migration of astrocyte precursors that form the dorsal lattice. In contrast, our data could also suggest that TBX3 may function within astrocytes, controlling their development and loss of *Tbx3* in retinal progenitor cells, which affects dorsal retinal ganglion cells, simply causes hyaloid artery persistence at P9 in the *Tbx3* cKO retina. The fact that we are observing abnormal vascular plexus and astrocyte formation in the dorsal retina, supports the hypothesis that the underlying defect is the loss of dorsal retinal ganglion cells. In any case, these mice provide a fruitful new model to study retinal hypovascularization.

Patients with syndromes, including Norrie disease (ND) or Familial Exudative Vitreoretinopathy (FEVR), develop hypovascular retinas that lead to congenital blindness. ^48^ These diseases have been associated with mutations in Wnt/Norrie signaling pathway ligands (*LRP5, NDP*), receptor (FZ4) and a co-receptor (*TSPAN12*), as well as a kinesin family member 11 (*KIF11*). ^6–9^ Mutations found in these proteins have been studied using mouse models of these mutant alleles. ^48–53^ Canonical Wnt signaling occurs when Wnt ligands bind to their receptors, which leads to stabilizing β-catenin that can regulate target downstream genes. While most of these factors work in the canonical Wnt/Norrie signaling pathway, *KIF11* has been shown to function independently from β-catenin signaling in retinal angiogenesis. ^48^ In separate studies, TBX3 has been shown to work in Wnt signaling and with a kinesin protein. In pull-down assays, TBX3 has direct interaction with another kinesin family member, KIF7, and affects its function. ^24^ In vivo, TBX3 has been shown to work with, or downstream of, the Wnt signaling pathway in ureter, lung and human colorectal cancer. ^23,26,54^ In future studies, we plan to determine in which molecular pathway Tbx3 functions.

Another characteristic, besides hypovascularization, shared among some FEVR patients is a persistent hyaloid artery. ^55–59^ In mice, this defect has also been found with the loss of melanopsin (*OPN4*), however, in these mutant mice, the retina is hypervascular, rather than hypovascular. ^11^ In contrast, the *Tbx3* cKO has a persistent hyaloid artery as well as a hypovascular retina. Along with FEVR, these phenotypes are also present in a mouse model of Retinopathy of Prematurity, in which neonatal pups are exposed to elevated oxygen levels from P0 to P4. ^60^ In these mice, angiogenesis is globally affected, where only about 25% of the entire retinal surface was covered by P8. ^60^ In comparison, vessels covered approximately 25% of the dorsal *Tbx3* cKO retina. Therefore, it is possible that Tbx3 is required for the hypoxia signaling pathway either in neurons or in astrocytes. Moreover, the similarities between the phenotypic changes we observe in the conditional *Tbx3* knockout mouse and those in FEVR/ROP patients suggests the exciting possibility that the *Tbx3* cKO may serve as a new mouse model for these human diseases.

We discovered that TBX3-GFP is expressed in astrocyte precursors. We found that *Tbx3* loss causes a reduction in astrocyte precursors in the newborn retina. Astrocytes arise from the optic disc progenitor zone which surrounds the optic nerve disc. ^42,61^ During migration into the retina, they proliferate prior to contact with endothelial cells, since the endothelial cells cause their differentiation. ^61^ Our results could be explained by either a reduction in astrocyte precursor proliferation or reduced migration due to corrupt cues from the neural retina lacking *Tbx3*. Retinal ganglion cells release platelet-derived growth factor A-chain (PDGF-A), which attracts astrocytes, but PDGF-A also causes proliferation of these cells. ^62^ As we have also observed a loss of RGCs in the dorsal retina, it is possible that loss of *Tbx3* indirectly affects astrocyte proliferation/migration. It is also possible that in astrocyte precursors TBX3 contributes to the regulation of a gene that controls astrocyte proliferation - like the transcription factor Tailless (*Tlx*). ^63,64^ We previously found that misexpression of frog *tbx3* (also known as ET for ‘Eye T-box’) induced *tll* expression. ^15^ It will be important to investigate the role of TBX3 in regulating *Tlx* expression and astrocyte proliferation and/or migration.

Later, in the differentiated retina we found truncation of the astrocytic lattice in the dorsal-temporal region. Blood vessels were growing into the area, but they failed to form a superficial plexus. Endothelial cells build the superficial plexus in response to the chemical and structural cues of the underlying astrocytes. ^65–69^ Therefore, it is logical that loss of dorsal astrocytes due to loss of *Tbx3* would lead to superficial plexus loss in that region. Finding dorsal intermediate blood vessels in the absence of the superficial and sometimes deep layers is curious. The deep plexus formation has been linked to spontaneous activity from cholinergic starburst amacrine cells. ^70^ Meanwhile, formation of the superficial and deep, but not intermediate, plexus layers requires the activity of the hypoxic response pathway (via the von Hippel-Lindau factor (Vhl) found in amacrine and horizontal cells. ^71^ In a recent study that used single cell PCR to generate a genetic regulatory network in the mouse retina, *Tbx3* was found in amacrine and Müller glia. ^72^ Overexpression of TBX3 in retinal progenitors at P0 led to an increase of amacrine, retinal ganglion cells, Müller glia, and bipolar cells, but repressed formation of rod photoreceptors. ^72^ Therefore, in future studies, we will determine the effect of *Tbx3* loss on all types of retinal neurons.

## Acknowledgments

We would like to thank Jason Shandler, Robertha Barnes, Amanda Bielecki, Abigail Snow, Stephen Sanabria and Dominica Colavito for their excellent technical assistance. We would also like to thank Dr. Arvydas Matiukas for his assistance in using the confocal microscope in Neuroscience Microscopy Core. The anti-Islet1/2 antibody was obtained from the Developmental Studies Hybridoma Bank, created by the NICHD of the NIH and maintained at The University of Iowa, Department of Biology, Iowa City, IA 52242.

## Supplemental Figures, Tables & Movies

See attached file Supplemental Materials file

## Notes

**Funding:** This study was supported by the National Institutes of Health (R21 EY029114 to A.S.V; R01 EY012676 to W.J.B; R21 EY030654 to M.E.Z; R01 EY024373 to S.F.), and grants to the Ophthalmology and Visual Sciences Department from Research to Prevent Blindness (Unrestricted Award) and the District 20-Y Lions Club.

### Competing Interest Statement

The authors have declared no competing interest.

### Summary of Updates

In this new version, we have edited the abstract, reduced the text length and added supplemental data files. We have also added data showing endothelial cell staining with our astrocyte markers in cKO retinas (Figure 5) to clarify the interaction between these two cell types. Author Dr. Sabine Fuhrmann, who was seminal in establishing this project, was also added.

