## Supplemental Material for "Loss of Tbx3 in mouse eye causes retinal angiogenesis defects reminiscent of human disease"

**Supplemental Figure S1**

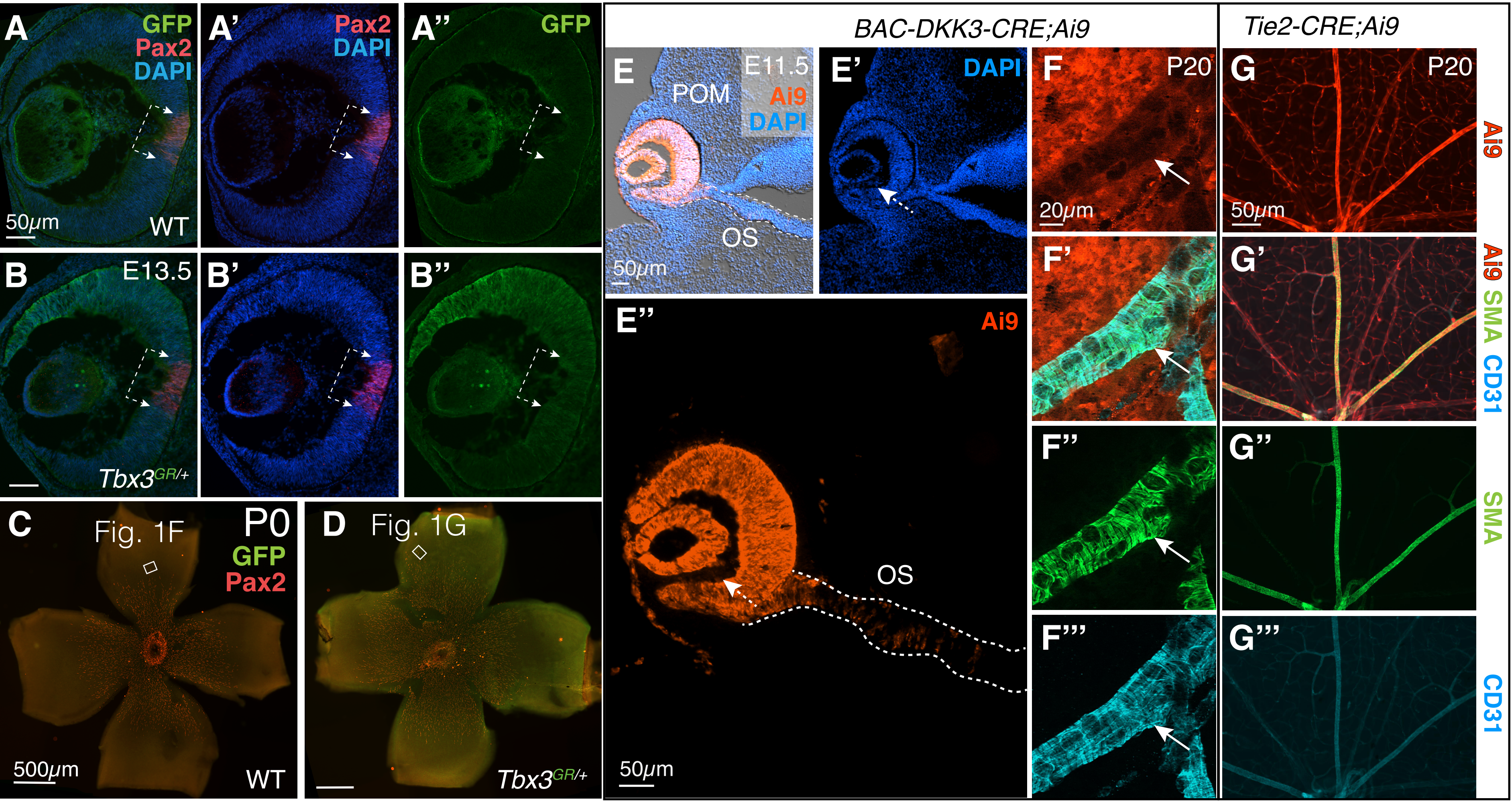


**Supplemental Figure S1. Tbx3 is expressed in embryonic neural retinal progenitors and Pax2-positive embryonic and perinatal astrocyte precursors.**

(A-B'') Embryonic retinal sections from sibling WT and Tbx3-GFP knock-in mice (*Tbx3$^{GR/+}$*) immunostained with anti-GFP (green) and Pax2 (red) antibodies (bracket marks Pax2-positive cells). DAPI labels nuclei. Scale bars indicate 50 μm. (C,D) Flat mounts from sibling P0 WT and *Tbx3$^{GR/+}$* immunostained with anti-GFP (green) and Pax2 (red) antibodies. Images were taken and enhanced using the same conditions. Box marks region imaged by the confocal microscope in Figs. 1F and 1G. Dorsal (D) is up in all panels. (E-E'') Ai9 Cre reporter crossed with *BAC-Dkk3-CRE* (*BAC-Dkk3-CRE;Ai9*) at E13.5. The only structure expressing tdTomato (red) is the neural retina, overlying ectoderm and optic stalk (OS). The red channel was enhanced to show that cells between the neural retina and lens (arrow) were not stained. (F-F''') At P20, a close-up of a blood vessel (arrow) stained with an antibody against alpha-smooth muscle actin (SMA) and blood vessel marker, CD31, shows a lack of tdTomato. Surrounding cells are expressing the Cre reporter. (G-G''') Another transgenic line expressing Cre under the control of Tie2 has overlapping expression with blood vessel markers,SMA and CD31.

### Supplemental Figure S2

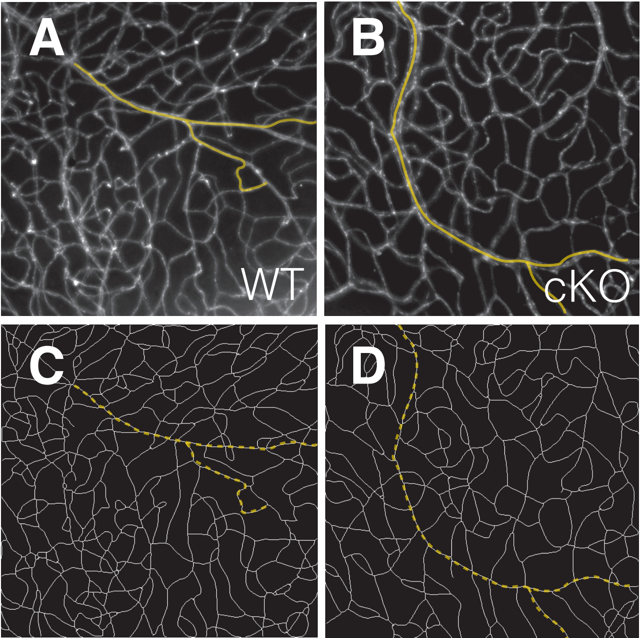


**Supplemental Figure S2. Example images of how branch points and density were calculated.**
(A,B) Flat mount retinas of CD31-positive dorsal retinal vasculature in WT and Tbx3 cKO mice; images were processed using ImageJ and (C,D) transformed into skeletonized versions of the corresponding images. The skeletonized images from WT (n = 3) and cKO (n = 3) were used for analysis. Yellow and dashed lines trace the same vessels in A and C, B and D.

### Supplemental Figure S3

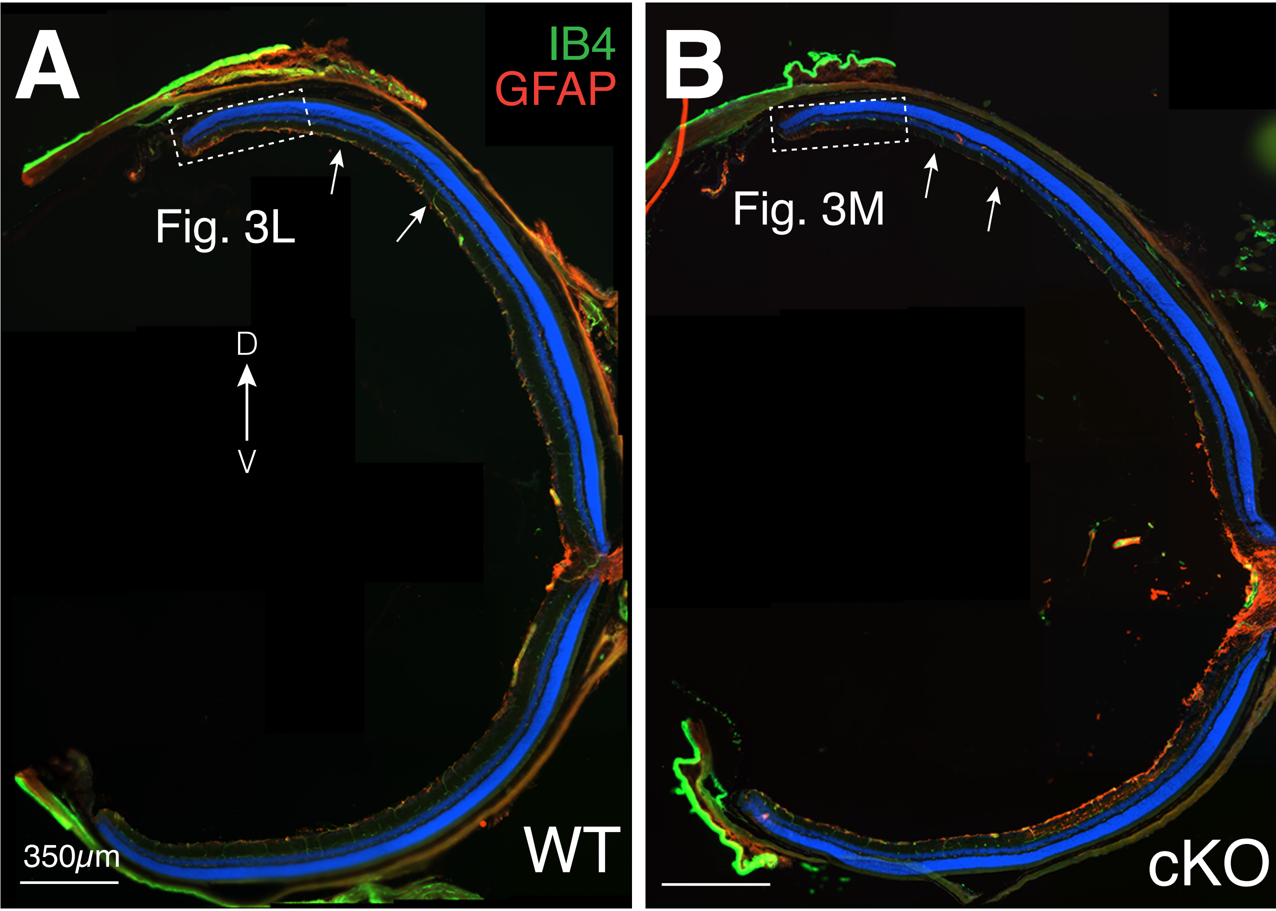


**Supplemental Figure S3. Blood vessel and astrocyte lattice formation are perturbed in P26 conditional knockout mice.**
A,B) Transverse sections (20μm) of P26 eyes through the optic nerve shows the regions taken to make Fig. 3 (dashed box). Blood vessels were stained with IB4 (green); astrocytes and Müller glia were stained with anti-GFAP antibody (red). Dorsal is up. Note the astrocytes in the dorsal inner limiting membrane in the wild-type (A, arrows), but much fewer in the cKO (B).

### Supplemental Tables

Supplemental Table S1. PCR primers and conditions for PCR reactions.

| Target | Primer name | 5'→3' Sequence | PCR conditions | | | | | |
|---|---|---|---|---|---|---|---|---|
| | | | Step 1 | Step 2 | Step 3 | Step 4 | Step 5 | Step 6 |
| **Tbx3-flox** | Tbx3 Geno01 | AAGTCATGGAGCTCGTATCGCG | 94˚ C 2 min | 94˚ C 20 sec | Repeat | 72˚ C 5 min | | |
| | Tbx3 Geno02 | GTGTGAGACAGAGAAATCAGTGG | | 55˚ C 20 sec | Step 2 | | | |
| | Tbx3 Geno03 | CCAACTGGTATCTTGATAAACCTC | | 72˚ C 20 sec | 35X | | | |
| **CRE** | 5' CreR 368 | AAAACGTTGATGCCGGTGAA | 94˚ C 2 min | 94˚ C 15 sec | Repeat | 72˚ C 5 min | | |
| | 3' CreR 922 | CCGGTATTGAAACTCCAGCG | | 60˚ C 15 sec | Step 2 | | | |
| | | | | 72˚ C 30 sec | 37X | | | |
| **Sex** | SX_F | GATGATTTGAGTGGAAATGTGAGGTA | 94˚ C 3 min | 94˚ C 20 sec | Repeat | 72˚ C 5 min | | |
| | SX_R | CTTATGTTTATAGGCATGCACCATGTA | | 57˚ C 20 sec | Step 2 | | | |
| | | | | 72˚C 20 sec | 35X | | | |
| **Tbx3$^{GR}$** | SH65A Tbx3-G | CATACGTGTATATGATGGGAGGTTG | 94˚ C 2 min | 94˚ C 20 sec | Repeat | 94˚ C 15 sec | Repeat | 72˚ C 2 min |
| | SH57A Tbx3-G | GCAATCTATACATGTCCTGCGAG | | 65-0.5˚ C 15 sec | Step 2 | 60˚ C 15 sec | Step 4 | |
| | | | | 68˚ C 10 sec | 10X | 72˚ C 10 sec | 28X | |
| **Ai9** | 5 Ai9WT oIMR9020 | AAGGGAGCTGCAGTGGAGTA | 94˚ C 2 min | 94˚ C 20 sec | Repeat | 94˚ C 15 sec | Repeat | 72˚ C 2 min |
| | 3 Ai9 WT oIMR9021 | CCGAAAATCTGTGGGAAGTC | | 65-0.5˚ C 15 sec | Step 2 | 60˚ C 15 sec | Step 4 | |
| | 5 Ai9WPRE oIMR9103 | GGCATTAAAGCAGCGTATCC | | 68˚ C 10 sec | 10X | 72˚ C 10 sec | 28X | |
| | 3 Ai9 oIMR9105 | CTGTTCCTGTACGGCATGG | | | | | | |

Supplemental Table S2. Antibodies and concentrations used for immunostaining.

| | Species raised | Company | Catalog # | Lot # | Block conditions | 1˚ Antibody concentration | 2˚ Ab Co. & cat# | Antibody ID | (PMID) |
|---|---|---|---|---|---|---|---|---|---|
| **GFP** | Chicken | AbCam | AB13970 | GR89472-2 2 | 1X PBS, 10% HIGS, 0.3% Triton | 1:1500 | AbCam AB150169 | AB_300798 | 33774011 |
| **Pax2** | Rabbit | Biolegend | 901002 (Covance# PRB-276P) | B287354 | 1X PBS, 10% HIGS, 0.3% Triton | 1:1000 | Invitrogen A21428 | AB_2565001 AB_291611 | 19505455 33428890 |
| **GFAP** | Mouse | Sigma | G3893 | 122914 | 1X PBS, 5% HIGS, 0.3% Triton | 1:500 | Invitrogen A21424 | AB_477010 | 6198232 26996101 |
| **PDGFRα (CD140a)** | Rat | BD Biosciences | 558774 | 8109820 | 1X PBS, 5% HIGS, 5% HIDS, 0.3% Triton | 1:500 | Jackson 112-605-167 | AB_397117 | 28943241 8982160 8875964 |
| **OPN4** | Rabbit | Advanced Targeting Systems Bio | AB-N38 (clone UF006) | 135-4 | 1X PBS, 5% HIGS, 5% HIDS, 0.3% Triton | 1:2500 | Invitrogen A31572 | AB_1608077 | 20503419 11823848 11834834 |
| **Islet1/2** | Mouse | DSHB | 39.4D5 | n/a | 1X PBS, 10% HIDS, 0.3% Triton | 1:100 | Invitrogen A31570 | AB_2314683 | 1350865 31313880 |
| **CD31** | Rat | BD Biosciences | 550274 (clone MEC13.3) | 8079850 | 1X PBS, 5% HIGS, 0.1% Triton | 1:250 | Invitrogen A11077 | AB_393571 | 9284815 7956830 |
| **IB4-FITC** | Griffonia simplicifolia | Sigma | L2895 | 125M4177V | 1X PBS, 5% HIGS, 0.3% Triton | 1:50 | - | - | 7107706 29511172 |
| **αSMA-FITC** | Mouse | Sigma | F3777 (Clone 1A4) | n/a | 1X PBS, 5% HIGS, 0.3% Triton | 1:500 | - | - | 29874128 |
